# Genome size variation is attributed to adaptive purging of transposable elements

**DOI:** 10.1101/2025.09.26.678839

**Authors:** Evgenii Potapenko, Bar Shermeister, Tali Mandel, Sariel Hübner

## Abstract

A substantial variation in genome size has been observed among individuals of the same species. Theory predicts that increased genome size may confer an advantage in populations with small effective size. However, contradictory evidence for the correlation with environmental variation, and limited understanding of the underlying genetic mechanisms, has cast doubts on the adaptive role of genome size variation. To address this, we studied two *Hordeum* species which were collected at the same sites along a wide range of environments but different in life habit (annual/perrenial) and mating strategy (self/out-crossing).

We observed significant variation in genome size, with differences of up to 10% consistently detected in both species. Mating system influenced only the proportion of variation maintained within populations but not the overall range of genome size. In both species, drought emerged as the primary environmental factor associated with genome expansion, with transposable elements (TE) accumulation identified as the main driver of this expansion. Genome-wide association studies revealed that TE silencing is the key mechanism regulating genome size, and that selection favors smaller genomes among individuals growing at suitable habitats. Under increasingly stressful conditions, the regulation of TE activity fades, leading to TE accumulation and genome expansion, thereby increasing genetic variation available for selection. This integrative study provides a comprehensive view of how genome size is regulated in natural populations and highlights its exaptive role in inducing genetic variation under environmental stress.

## Introduction

The DNA content of an organism, known as the C-value, varies extensively among taxa, with a wide and gradual increase in genome size with the species complexity (Schubert et al., 2016). Despite this clear trend along the tree of life, the correlation between genome size and complexity is faded at the sub-kingdom resolution where increased genome size does not seem to provide additional functional features or complexity (Mirsky & Ris, 1951). This perplexing phenomena of DNA accumulation with no clear functional gain was named the “C-value paradox” (Thomas 1971), and later it became clear that organisms contain much more DNA than required to encode proteins and regulatory sequences (Cavalier-Smith, 1985). In practice, some organisms contain massive genomes which pose a significant burden during the cell cycle (Bennett & Leitch, 2005; Lynch & Conery, 2003). With the discovery of repetitive elements and their volume in the genome, a new term was proposed, “the C-value enigma”, to capture the unclear biological purpose of non-coding DNA accumulation (Gregory, 2001). Over the years, the contribution of non-coding DNA to regulation of gene expression, epigenetics and other processes became clearer, however the question of non-coding sequence obesity despite its clear biological burden remains largely elusive.

To elucidate the genetic mechanisms that regulate genome size variation, research has focused on intraspecific variation where taxonomic differences, ploidy, mating strategy and other factors are controlled. Studies in maize (Rayburn & Auger, 1990), sunflower (Michaelson et al., 1991), soybean (Graham et al., 1994; Rayburn et al., 1997), pea (Arumuganathan & Earle, 1991) and other species, were able to find correlations with environmental gradients including temperature, altitude and latitude (Knight et al., 2005). Moreover, correlations between genome size variation, growth and morphological features were highlighted in the context of cell size (Herben et al., 2012), leaf mass per area (Morgan & Westoby, 2005) and minimum seed mass (Rayburn et al., 1997). These correlations were considered as evidence for an adaptive role of genome size variation along environmental clines (Benor et al., 2011; Du et al., 2017; Gallagher et al., 2011; Khorami et al., 2018; Krahulcová et al., 2017). In contrast, other studies have failed to identify significant correlations with either vegetative or reproductive traits, nor with temperature and precipitation gradients (Basak et al., 2019; Realini et al., 2016). The main arguments posed by the opponents of the adaptive hypothesis were that the observed correlations between environment, phenotypes and genome size variation are overinterpreted and underrate the effect of genetic drift and phylogenetic and mating system (Koonin, 2016; Lynch & Conery, 2003; Petrov, 2001; Whitney et al., 2010). Moreover, larger genome increases the probability for lethal mutations (LaBar & Adami, 2020), and requires higher energy resources (Knight et al., 2005). Despite the central role of genome size variation in evolution and speciation, it is unclear whether genome size variation is an outcome of neutral demographic processes associated with maintenance of diversity (Lynch & Conery, 2003) or a mechanism to promote adaptation under changing environments (Jordan et al., 2015; Mei et al., 2018). Thus, the adaptive role of genome size variation under stressful environmental conditions remains controversial.

The era of whole genome sequencing made it clear that gene content has little contribution to genome size variation. In plants, the main factors determining the size of the genome are abundance of spliceosomal introns, repetitive sequence, and specifically transposable elements (TEs) (Lynch & Conery, 2003; Petrov, 2001). TEs exhibit broad diversity in their structure and transposition mechanisms (Feschotte & Pritham, 2007), where retrotransposons play a major role in genome augmentation. The most abundant retrotransposons in plants are the long terminal repeats (LTR) which were found to be responsible for large and rapid genome size increases in *Panax ginseng* (Lee et al., 2017), *Oryza australiensis (Piegu et al., 2006)* and *Oryza granulata* (Ammiraju et al., 2007). The activation of TEs is responsible not only for increased genome size but also as agents of novel genetic diversity for rapid adaptation to environmental change (Stapley et al., 2015).

The genus *Hordeum L.* is part of the *Triticeae* family and include over thirty species that are widespread in the northern hemisphere, in South Africa and South America (Bothmer et al., 1995). This genus include species characterized by different life cycles (annual vs. perennial), mating systems (self-crossing vs. out-crossing), and ploidy levels (2n, 4n, 6n) (Bothmer et al., 1995). Moreover, *Hordeum* species are characterized with large genomes and high TEs content, occupying circa 80% of the genome, mainly LTRs (Wicker et al., 2017). Among the species in the genus, barley (*H. vulgare*) and its wild progenitor *H. spontaneum* are the most studied due to their economical and agricultural importance (von Bothmer & Jacobsen, 2015). The wild progenitor of cultivated barley, *H. spontaneum* (HS) has a 5Gbp diploid genome (2n = 14), is a self-crossing species, and is highly abundant along the Fertile Crescent (Bothmer et al., 1995; Monat et al., 2019). A sister species to wild barley is *H. bulbosum* (HB) which has a diploid cytotype (2n = 14) that occurs west to Greece and a tetraploid cytotype that occurs to the east (2n = 28) (Blattner, 2006; Feng et al., 2025; Jörgensen, 1982). It is an obligatory out-crossing species with a perennial growth habit ought to its ability to form bulbs at the basis of each tiller (Fuerst et al., 2023). The two species were split phylogenetically 4M years ago (Blattner, 2018) and share many taxonomic, ecological and morphological characteristics despite the substantial differences between them. Therefore, the HS-HB system provides a good model for studying the regulation of genome size variation.

To elucidate the main factors regulating genome size variation, we established a comprehensive dataset for two germplasm collections for HS and HB, both collected at the same sites along a wide environmental gradient. These collections provide a comprehensive and comparative platform to investigate the effect of ecological, demographic, genetic, and evolutionary factors on intraspecific genome size variation. Our results highlight the main environmental factors associated with genome size variation across species and the genetic factors regulating genome size variation.

## Results

### Mating strategy effect genome size variation within populations while the range remains consistent

Among the 31 species in the genus *Hordeum*, *H. spontaneum* (HS) and *H. bulbosum* (HB) are particularly abundant in Israel, where they grow in large stands across diverse habitats. We sampled populations at 30 different locations spanning a wide eco-geographical range, from the Mediterranean timberline zone to the arid regions near the Dead Sea. Among sampling sites, HB was absent from several sites mainly in the desert, thus both species co-occur at 22 locations (Fig. 1a,b). The sampling locations varied widely in key environmental factors, including maximum temperatures during the flowering season in March (ranging from 11–24 °C), average annual precipitation (248–1,184 mm), and elevation (-204-1,445 meters above sea level).

**Figure 1.**
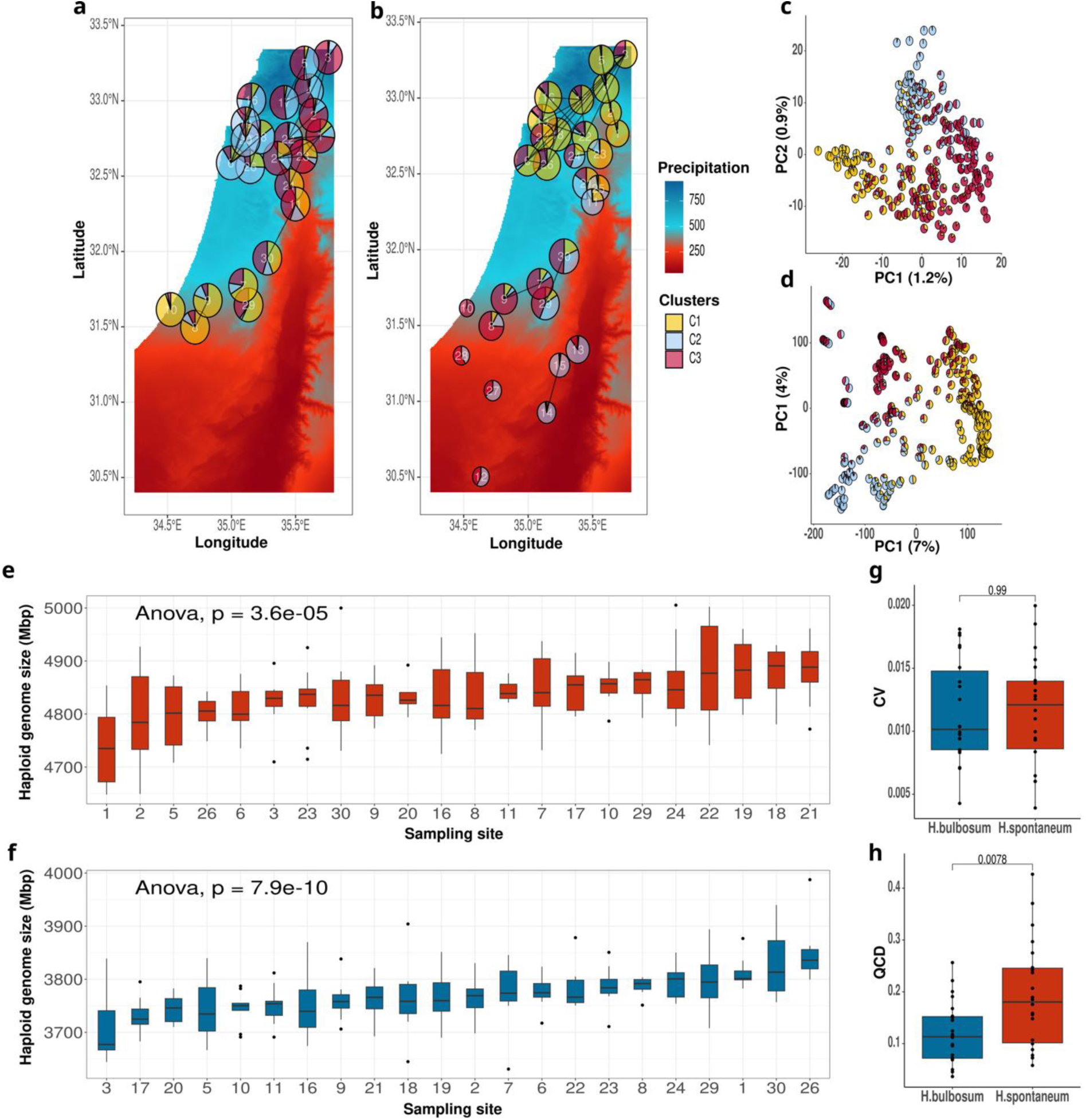
Population structure and genome size variation in *H. spontaneum* (HS) and *H. bulbosum* (HB). Maps showing sampling locations of HB **(a)** and HS **(b)** populations over a gradient of annual precipitation (dry = red). Pie charts represent the proportion of assignment to genetic clusters in each site, and the pie size reflects average nucleotide diversity. Black lines indicate connections between populations with inferred gene flow. Sampling site identifiers are indicated inside pie charts. Principal component analysis for HB **(c)**, and HS **(d)** with samples colored according to their assignment to genetic cluster. Genome size variation across populations of HS **(e)**, and HB **(f)**. In the boxplots, sites where both species co-occur are presented, *p*-value from ANOVA test are indicated at the top of each plot. Comparison of genome size coefficients of variation between species **(g)**, and quartile coefficients of dispersion **(h)**, where *p*-values obtained from Wilcoxon test are indicated at the top of each boxplot.

Whole genome sequence data for the entire HS collection was processed and 33M variants were called after stringent filtering (Potapenko et al., 2025). For HB, genotyping-by-sequencing (GBS) data was generated for the entire collection and reads were aligned to the haplotype resolved tetraploid reference genome (Feng et al., 2025), thus a total of 16,776 SNPs were called after filtering.

To investigate population stratification and gene flow in each species, both SNP datasets were filtered to reduce statistical bias by pruning markers in linkage disequilibrium (LD > 0.2). Overall, populations of both species clustered according to the three ecotypes previously identified in HS (north, coast, and desert; (Hübner et al., 2009, 2013), with clear clinal variation observed between ecological clusters (Fig. 1a-d). As expected, stronger population divergence and stratification were observed in the predominantly self-crossing HS, compared to the outcrossing species HB, which exhibited higher levels of gene flow across greater distances (Fig. 1a,b). Population structure analyses were further supported by an AMOVA, which revealed a striking difference between species. In the outcrossing HB, 89% of the genetic variation was found within populations, whereas in the self-crossing HS, only 40% of the variation occurred within populations (Fig. S1). Although the analyses in each species were conducted with a different type of genotyping data the differences are substantial and attributed to mating strategy. To assess the relative effect of geographic versus environmental distance on genetic variation, a partial Mantel test was performed. The results indicated that population divergence in HS was much more influenced by geographic distance than in HB (Fig. S2). Together, these analyses support higher admixture in HB compared to the self-crossing HS.

Next, genome size was measured for all individuals across species using flow cytometry. To enable a direct comparison of genome size variation between species, HS accessions collected from sites where HB was absent were excluded from the analysis. A substantial variation in genome size was observed, ranging from 3.7 to 4.1 pg for haploid genome in HB, and 4.7 to 5.1 pg for HS (Fig. 1e,f). Interestingly, despite the considerable differences between the two species in ploidy, mating system, and life habit, the total variation in genome size was 8-10% in both species, implying a biological limitation for the extent of genome size variation within a species. This was further supported by a comparison of the coefficient of variation (CV) in genome size across populations designating similar overall range in HS and HB (t = -0.28, df = 46.4, p = 0.78; Fig. 1g). Analysis of the proportion of within-population genome size variation in each species using the quartile coefficient of dispersion (QCD) revealed significantly higher variation within populations in the self-crossing HS compared to the outcrossing HB (t = 2.48, df = 47.83, p = 0.0077; Fig. 2h). These results are consistent with the AMOVA findings, suggesting that the higher gene flow in the outcrossing species has a homogenizing effect, whereas lower gene flow in HS maintains substantial differences in genome size within the same population.

**Figure 2.**
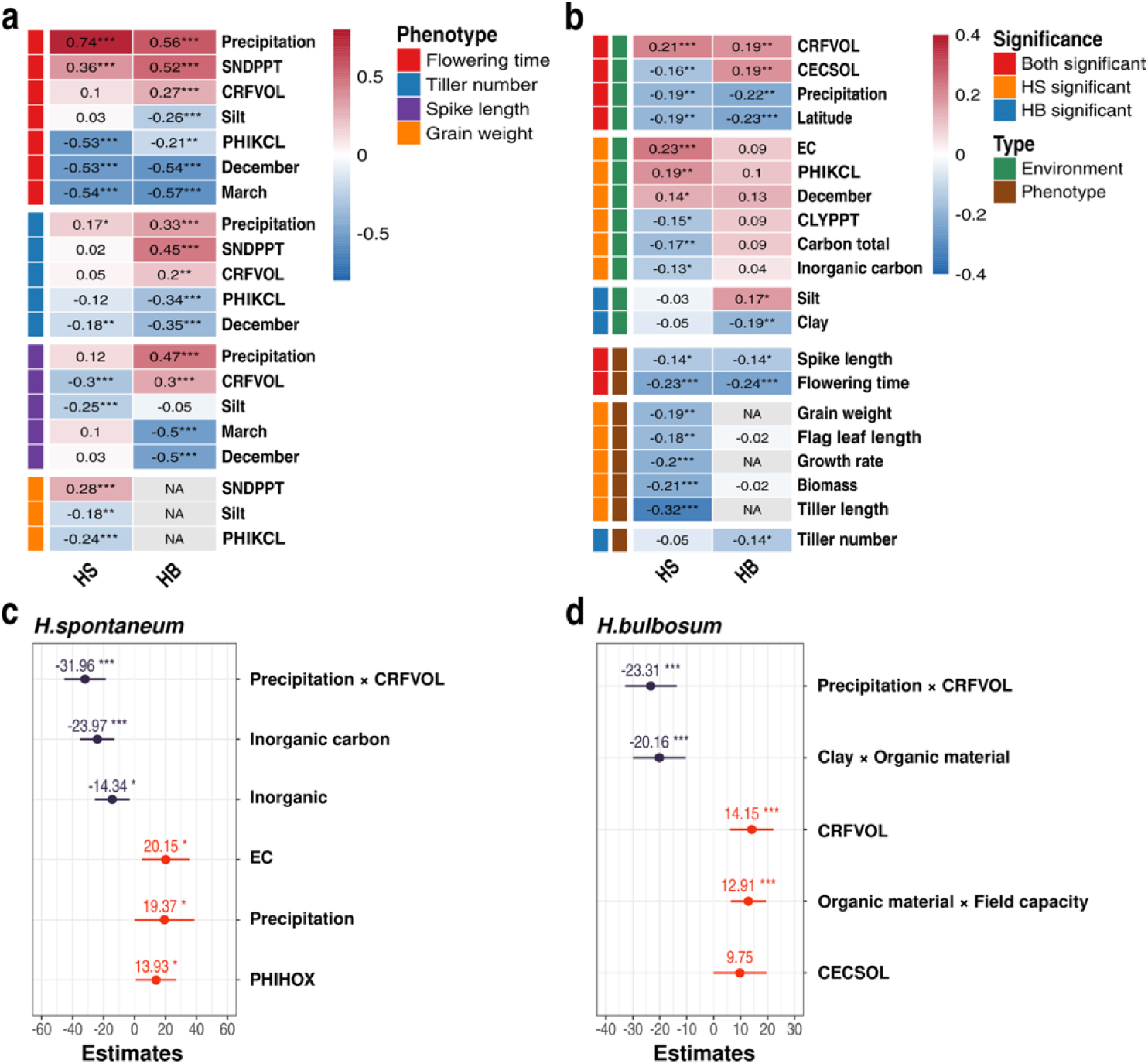
Association between genome size, phenotypes and environmental gradients. Heatmap for the correlations between environmental factors and phenotypic traits **(a)**, with strongest associations highlighted. Heatmap for the correlations between genome size with environmental factors, and with phenotypic traits **(b)**. Fixed effect estimates from linear mixed-effects models (LMER) for genome size variation in HS **(c)** and HB **(d)**, with sampling site included as a random effect. Error bars represent 95% confidence intervals. Asterisks indicate significance levels: *** P < 0.001, ** P < 0.01, * P < 0.05.

Nevertheless, the overall variation in genome size across all populations is consistent between species.

### Genome expansion is correlated with environmental stress

To evaluate the association between environmental stress, genome size and adaptation, we first compared the variation in phenotypic traits along environmental gradients in HS and HB. To provide a proper basis for comparison between species, we focused on individuals collected at the 22 sites where both species co-occur. Climatic data for each sampling location were retrieved from the Israeli Meteorological Service using sampling site-specific coordinates (Table S1). Soil data was obtained from the ISRIC database with 1km resolution (Hengl et al., 2014) and from lab measurements of soil samples taken at each site during the collection excursions (Table S1). Plants were grown in pots and full irrigation under common garden conditions in a randomized block design, and phenotypes were collected along the season (Table S2).

Flowering time was strongly influenced by water availability in both species, with plants flowering earlier under drier conditions. This effect was stronger in HS (r = 0.74; Fig. 2a) than in HB (r = 0.56; Fig. 2a, Fig. S3, Table S3). In HB, flowering appeared more sensitive to the volume of coarse fragments (CRFVOL), suggesting a stronger effect of the soil composition in plants which rely on bulbs for propagation (r_HS_ = 0.10, r_HB_ = 0.27, Fig 2b). Flowering time was also strongly associated with average temperature equally in both species (r_HS_ = -0.53, r_HB_ = -0.55 for average minimum temperatures in December) reflecting earlier flowering under warmer conditions. In HB, the tillers number was significantly correlated with environmental clines, where individuals from cooler temperatures and higher precipitation tend to produce more tillers. In HB, a bulb propagation organ is developing at the basis of each tiller, thus more tillers indicate higher propagation capacity. In contrast, tiller number tends to be a plastic trait in HS with no clear correlation with environmental gradient (Fig 2b). Grain weight was measured only in HS and was positively correlated with the soil fractions composition and pH, highlighting the importance of edaphic heterogeneity in shaping key traits. Together, these findings demonstrate distinct environmental correlations of key fitness-related traits in HS and HB, reflecting their contrasting ecological strategies as an annual and self-crossing versus a perennial and outcrossing species.

Next, we examined the correlations between environmental variables and genome size variation, leveraging samples from all sites to maximize statistical power, rather than restricting the analysis to only common sampling locations. Both species showed negative correlations between genome size and precipitation, and positive correlations with the soil coarse fragment volume (*p* < 0.001, Fig 2b, Table S4). These trends support the hypothesis that genome expands under stressful conditions, whereas the strongest stress is imposed by shortage of accessible water. Additional significant correlations of soil electrical conductivity (EC) and pH (PHIKCL) were observed only in HS which reflects its higher occurrence in the desert where soil salinity stress is also a limiting factor. In HB, genome size correlated positively with silt and negatively with clay, which reflects the preference of bulbing plants in well-drained soils.

To refine these results and estimate the relative contribution of environmental factors, we implemented a mixed-effects linear model for each species separately using available data from all populations of HS and HB. For HS, the model accounted for 29.7% of total genome size variation, with environmental predictors explaining 22.3% and the remaining variation is attributed to differences between sampling sites (Fig. 2c, S4a). For HB, the entire model explained 31.4% of the variation, and the environmental predictor alone explained 27.6% (Fig. 2d, S4b). Thus, indicating that most spatial variance in both species is captured by environmental factors rather than merely geographic distance. The strongest predictor of genome size in both species was the interaction between precipitation and coarse fragment content, which showed a significant negative effect (HS: β = –31.96, p < 0.001; HB: β = –23.31, p < 0.001). This indicates that drought due to low precipitation and low water capacity in the soil are the dominant stressors and promote genome expansion in both species. In HS, higher electrical conductivity (β = 20.15, p = 0.017) was also significantly associated with genome size increase, suggesting that salinity stress, common in desert saline soils, is also a key driving factor.

Finally, we assessed correlations between genome size and the phenotypic traits measured for each species. In HS, nearly all observed traits, including tiller length, biomass, flowering time, grain weight, spike length, and flag leaf length were negatively correlated with genome size. In HB, only tiller number, spike length, and flowering time showed significant negative correlations (Fig. 2b, Table S4). This pattern indicates that larger genomes are associated with reduced biomass and accelerated flowering, especially in HS, supporting the association between stress and genome expansion.

Taken together, these findings show that both species exhibit genome size increases under drought-related stress, yet they diverge in their sensitivity to other environmental stressors. The distinct yet overlapping response patterns observed highlight the central effect of stress on fitness related traits and genome size variation regardless of the differences between species in mating strategy, life history and growth habit.

### Genome expansion is driven by transposable elements accumulation

In many higher plant species and specifically among grasses, a large fraction of the genome is comprised of repetitive sequences. In HS and HB, more than 80% of the genome corresponds to repetitive elements of which the long terminal repeats (LTRs) account for most of the repetitive fraction of the genome. To quantify the abundance of repetitive elements, we analyzed whole genome sequence data available only for HS accessions using a reference-free approach by performing a *de novo* assembly procedure for a representative fraction of reads and quantifying the abundance of repeats in the genome (Fig. S5). This procedure enabled us to obtain unbiased estimates for the proportion of repetitive sequences along the genome in each accession at a reasonable computational cost. We focused on the fraction of LTRs as the major dynamic portion of repetitive sequences in the genome, which was highly variable among accessions. Overall, the variation in LTRs content extended over 202.1Mbp, which accounts for 54.6% of the total variation in genome size estimated by the flow-cytometry. A Pearson’s correlation test between genome size and amount of LTRs indicated a strong significant concordance (r = 0.82, p < 2.2e-16), thus the proportion of LTRs is a major feature contributing to genome size variation in HS (Fig. 3a).

**Figure 3.**
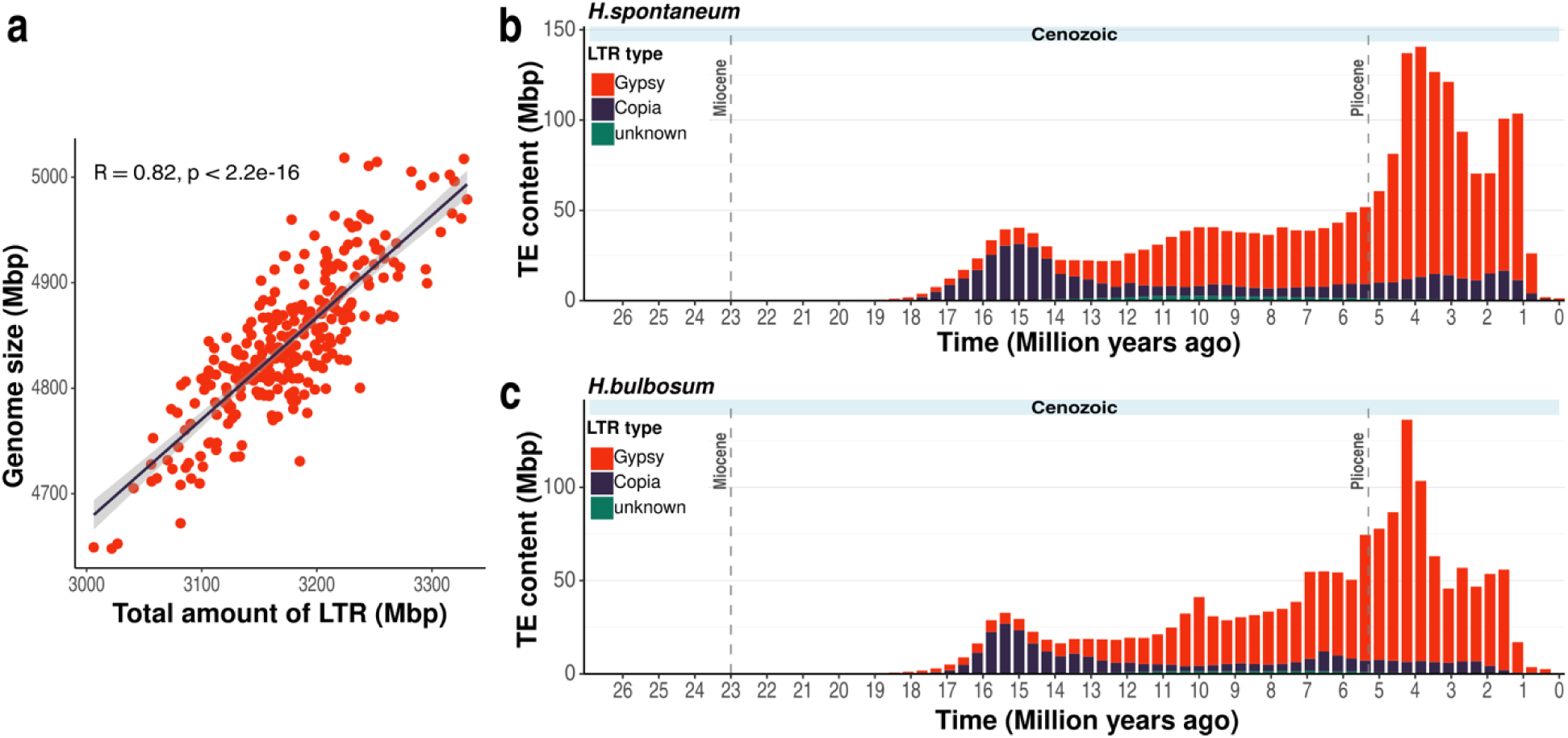
Transposable element dynamics in *Hordeum spontaneum* (HS) and *H. bulbosum* (HB). **a)** Correlation between genome size of HS and the total amount of LTR retrotransposons. Divergence landscapes of LTR retrotransposons in HS **(b)** and HB **(c)** with sequence divergence converted to million years ago.

TE accumulation frequently occurs in bursts of proliferation and can therefore lead to rapid genome expansion. To explore the course of TE accumulation, we calculated the sequence divergence among TEs families using the non-redundant TREP database (Wicker et al., 2002) and a reference genome of each species. As a representative genome for wild barley, we used the B1K-04-12 V1 assembly (Jayakodi et al., 2020) that was sampled also in the study region (Hübner et al., 2009). The degree of sequence divergence reflects the time elapsed since TEs proliferation, while their copy number provides an estimate for the magnitude of the amplification burst. The two species share a prominent peak for LTRs with circa 10-11% divergence, consistent with an amplification event coinciding with the split between HS and HB approximately 4M years ago (Fig. 3b,c). In HS, another recent LTRs burst event was identified which is absent in HB, with 3-4% divergence. In addition, we investigated the historical accumulation of non-LTR TEs (primarily DNA transposons) following the same procedure (Fig. S6). Interestingly, the observed TE burst in both species showed amplification burst with divergence around 15%. This difference between LTRs and non-LTRs may be attributed to a distinct mutation rate between class-I and class-II TEs, the effect of selective constraints between classes, or asynchronous transpositional activity in response to environmental activation.

### Stress alters selection against transposable elements activity

To further investigate the genetic regulation of genome size variation, we performed a genome-wide association study (GWAS) in each species separately, with the estimated genome size as a quantitative trait in the model, the genotype data as predictor and population structure as covariance. The high-resolution genotype data for HS enabled us to clearly identify a single genomic region which was strongly associated with genome size variation (Fig. 4a). This region is located on chromosome 3H and spans over 5.9 Mbp with well distinguished boundaries (3H:110,311,085–116,513,950, Fig. 4b). Interestingly, despite its considerable length, this region contains only 52 genes of which several are potentially attributed to transposon regulation and silencing (Table S5). For example, within this region we identified an *Argonaute* gene (ARO), a key component of the RNA-induced silencing complex (RISC), which plays a central role in suppressing TE activity in Arabidopsis and controlling genome expansion (Durán-Figueroa & Vielle-Calzada, 2010; Underwood & Martienssen, 2015). The ARO gene was also reported to regulate grain size through miRNA pathways in rice (Zhong et al., 2019), thus suggesting a pleiotropic effect of ARO. Adjacent to the ARO, we also identified a gene encoding a subunit 4 of the CCR4-NOT transcriptional complex (CNOT4). Recent studies in Arabidopsis have shown that CCR4 targets TE transcripts that escape siRNA-mediated regulation, thereby helping to control transposon mobilization (Wang et al., 2024). Other interesting genes found within this region are the ATP-dependent DNA helicase PIF1, which has been implicated with linking environmental signaling and TE silencing pathways (Ammari et al., 2024) and *ALP1-like* which is involved in repression of TEs loci in Arabidopsis (Liang et al., 2015).

**Figure 4.**
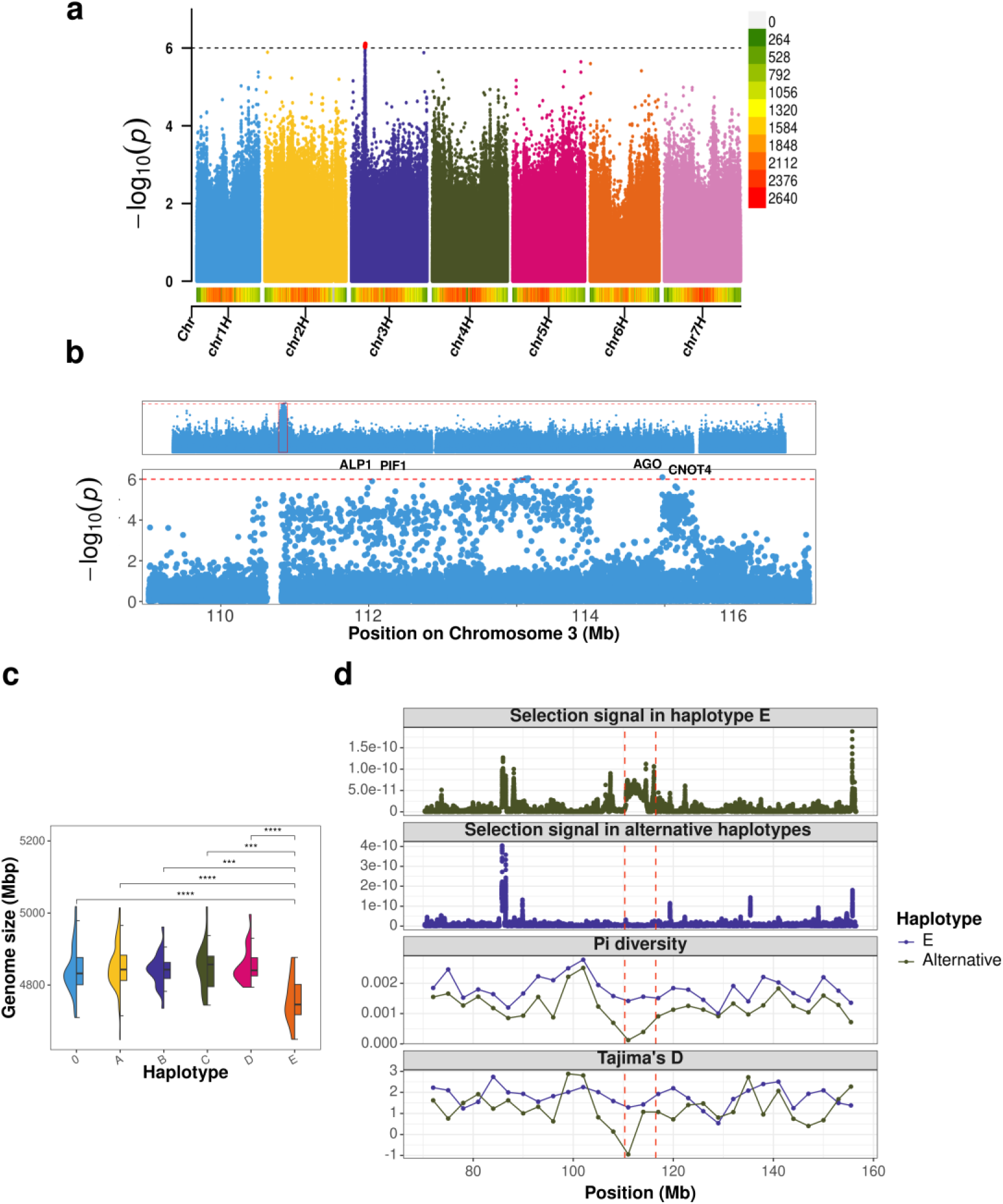
Genomic analysis of genome size variation. **a)** Genome-wide association of genome size variation in *H. spontaneum*. SNP density is shown in the colored bar at the bottom of the plot with gradient track from green to red (low-high density). **b)** Zoom-in Manhattan plot of the association signal with genome size variation. **c)** Boxplot showing genome size differences between individuals carrying different haplotypes (including samples shared no haplotypes labeled as 0 ), highlighting the haplotype-level effect. Asterisks indicate significance levels: **** P < 0.0001, *** P < 0.001, ** P < 0.01, * P < 0.05. **d)** Footprints of selection for haplotype E and other haplotypes in the identified genomic region (marked with red vertical dashed lines) with ±40 Mbp flanking context.

Performing GWAS in HB, has yielded a single signal on chromosome 4H_1 (Fig. S7). Due to the low genotyping resolution of GBS, we defined a larger region spanning 500Kbp around this locus to search for candidate genes. Similarly to the results for HS, this region had low gene density (Table S6). Among the 14 annotated genes found in this region, five were identified as transposon-annotated genes and the remaining nine genes included several intriguing candidates. For example, a ATP-dependent RNA helicase which could potentially be involved in eliminating transcripts produced by transposons (Wu-Scharf et al., 2000), a DDT-domain– containing gene which are involved transposable elements repression (Zhang et al., 2023), and a stress-response NST1-like protein which is involved in abiotic stress signaling (Mitsuda et al., 2005). Although most of the functional support is obtained from Arabidopsis, the association signal in both species highlights the involvement of TE repression and silencing in genome size variation.

To further validate the GWAS results, we tested for differences in genome size between individuals carrying alternative alleles at the identified loci. In HS, a large genomic region spanning 5.9Mbp was identified with a dense cluster of associated SNPs. Within this region, five haplotypes (A–E) were identified, with haplotype E exhibiting a significant reduction in genome size (Fig. 4c, S8). Notably, this haplotype was found across individuals from different sampling sites, indicating that genetic relatedness and demographic effects are unlikely to be the main drivers. In HB, only one SNP was significantly associated with genome size, likely due to the limited marker density of GBS. Haplotype-based analyses were not feasible; thus, genome size was compared between alternative alleles of the identified SNP. Individuals with the minor allele were also characterized with larger genomes, yet this finding is based on 10 individuals and should be further explored with higher resolution of genotype data (Fig. S9).

To test whether genome expansion is subject to selection and therefore potentially adaptive, we applied the RAiSD analysis to detect signatures of selective sweeps among individuals carrying the haplotype E and other haplotypes (Fig. 4e). Interestingly, a strong signal of selection was identified exclusively among individuals carrying the haplotype E, which is associated with a smaller genome. This finding was further supported by a substantial reduction in nucleotide diversity and negative Tajima’s D values, both consistent with footprints of selection (Fig 4e). In contrast, no evidence of a selective sweep was observed among individuals carrying other haplotypes which are characterized with larger genomes.

Notably, individuals carrying haplotype E originated from habitats which were previously identified as highly suitable for wild barley based on ecological niche models (Potapenko et al., 2025). Together, these results suggest that silencing of TEs, which contributes to maintenance of smaller genomes is under selection to preserve genome integrity in thriving individuals. In contrast, under stressful environmental conditions purging of TEs becomes less efficient, leading to genome expansion through accumulation of TEs.

## Discussion

We investigated two closely related *Hordeum* species that are abundant across a wide range of environments. These species co-occur in the same habitats, but differ in mating system, life history, and ploidy level, thus providing an ideal system to explore the evolutionary mechanisms underlying intraspecific genome size variation.

### Is intraspecific genome size variation biologically constrained?

Despite substantial variation between populations, both HS and HB showed a consistent genome size range of 8–10%. This range lays between previous estimates of circa 3% (Jakob et al., 2004) and 18% (Kankanpaa et al., 1996) for *Hordeum*, yet earlier estimates were made based on a limited sample sizes and were lacking representation of ecological diversity. The consistent 10% difference observed in this study across comprehensive diversity panels of two distinct species suggests a possible biological limitation to genome size variation within a species although these estimates may vary in other systems. Despite the consistency of genome size range in HS and HB, the pattern of variation differed between species. In the selfing species HS, greater within-population genome size variation was observed compared to the outcrossing HB. This is potentially due to the extensive gene flow in HB which homogenizes genome size differences at the local scale but allowing sufficient differences across broader geographic ranges. In contrast, in a selfing species which rarely hybridize, substantial genome size differences can accumulate at the local scale without significantly affecting population persistence. These findings elaborate previous comparative studies examining mating system effects on genome size evolution across a wide phylogenetic representation (Whitney et al., 2010). Here, we were able to identify a clear effect of mating system but only at the population level while the overall range remains consistent between HS and HB, thus highlighting the importance of microevolutionary scales in understanding the broader evolutionary process.

### How is environmental stress associated with genome size variation?

The unique germplasm collections used in this study which include two different species that co-occur at the same habitats along a wide range of environmental conditions offer a comprehensive and comparable system to address a long-standing question of how environmental stress influences genome size variation. Previous studies reported correlations between genome size and environmental factors including temperature and precipitation in Lilium (Du et al., 2017) and maize (Realini et al., 2016), altitude in maize (Bilinski et al., 2018) and in Dactylis glomerata (Reeves et al., 1998). Similar correlations have been observed among *Hordeum* species where temperature has been identified as the major factor (Turpeinen et al., 1999). Together, these studies suggest an adaptive role of larger genomes. Multiple, mechanisms have been proposed to link genome size variation and adaptation. In maize and teosinte, genome size has been inferred to covary with cell-production rate and earlier flowering under shorter growing season (Bilinski et al., 2018). Comparative analyses across angiosperms suggested that larger genomes are associated with increased guard-cell size and reduced stomatal density, potentially constraining developmental rates (Beaulieu et al., 2008). Broader sampling of vascular plants indicated that genome size predicts cell size and tissue packing densities, which may limit the maximum photosynthetic capacity and influence ecological strategies (Roddy et al., 2020). Positive associations were also reported between genome size and seed mass, pointing a potential link with dispersal and establishment (Beaulieu et al., 2007). Although these indications are largely correlative, they suggest a potential adaptive role for genome size variation.

We observed significant correlations between environmental gradients and genome size variation in both species, with precipitation and soil water-capacity emerging as the most influential factors (Fig. 2). Larger genomes were more frequent under harsher environmental conditions, with the largest genomes observed in desert isolated habitats (Table S4). This corroborates with the theory conferring that larger genomes provide a compensation for loss of diversity among populations with low *N_e_*due to demographic or selective constraints (Lynch & Conery, 2003). Individuals with larger genomes tended to flower earlier, grow more rapidly and maintain smaller seeds in both species, suggesting an escape strategy under harsh conditions rather than direct phenotypic effect of genome size as previously suggested (Beaulieu et al., 2007, 2008; Roddy et al., 2020). In accordance with other studies, the correlation coefficients in our study were significant but relatively low, indicating that environmental factors account for only a fraction of the observed genome size variation. These correlations may be interpreted as a positive contribution of larger genomes to adaptation (Beaulieu et al., 2007, 2008, Bilinski et al., 2018), however the link to an adaptive biological mechanism has been inconclusive.

### How can transposable elements dynamics contribute to adaptation?

Our analysis of transposable elements (TEs) was limited to HS due to the lack of whole-genome sequence data for HB, thus restricting the generalization of our findings. Nevertheless, several lines of evidence from the GWAS suggest that similar processes may occur in both species, supporting the relevance of our results also to other species.

In HS, TEs comprise approximately 80% of the genome, with long terminal repeats (LTRs) being the most abundant (Fig. S10). We showed that the proportion of LTRs is strongly correlated with genome size (r = 0.82, Fig. 3a), highlighting their central role in genome expansion. TEs are a major source of genetic variation and are known to proliferate rapidly under stressful conditions (Baduel et al., 2019; Bourque et al., 2018; Esnault et al., 2019), which aligns with the observed increase in TE content, and thus larger genome size in populations from stressful environments (Fig 2c,d, Fig. 3a). Interestingly, our GWAS, using genome size as a trait in both species, identified genes associated with TE purging and silencing (Fig. 4a,b, S7; Table S5, S6).

This suggests that genome size regulation primarily occurs through mechanisms that suppress TE activity, thereby preventing their accumulation. Moreover, we detected signs of selection around these loci, particularly in accessions with smaller genome, implying that TE silencing and controlled genome size is advantageous rather than TE accumulation and expansion (Fig. 4c,e). Therefore, larger genomes are indeed observed under more stressful environmental conditions, yet this pattern likely reflects ineffective TE silencing rather than adaptive genome expansion.

An efficient control of TEs activity enables to maintain high integrity of smaller genomes. Identification of a genomic region enriched for TE silencing genes provides a focal point for future functional studies to dissect the molecular mechanisms controlling TE proliferation and genome expansion and their interaction with environmental shifts.

Our study demonstrates that genome expansion under stress is not adaptive in itself; rather, it results from reduced regulation of TE activity. In well-adapted populations, flourishing under suitable conditions, selection favors effective TE silencing, thus helping to maintain high fitness and genome integrity. Under stress, these silencing mechanisms are deteriorated, leading to increased TE activity and genome expansion which is enhanced further in isolated populations in the desert. Despite their genomic burden, the exaptive accumulation of TEs rapidly generates new genetic variation, thereby increasing standing genetic variation available for selection (Koonin, 2016). As stress intensifies, TE proliferation increases accordingly, thereby maintaining a selection-mutation balance (Fig 3b,c).

## Materials and Methods

### Establishing the germplasm collections

Seeds of two wild barley species *Hordeum spontaneum* (HS) and *Hordeum bulbosum* (HB) were collected from sampling sites across Israel, representing various climatic and environmental conditions. Sampling sites were selected based on previous studies and a specific sampling design which enabled to maximize the representation of environmental and genetic diversity and minimize the confounding effect of demographic history and population structure (Hübner et al., 2009, 2012, 2013; Hübner & Kantar, 2021; Potapenko et al., 2025). Both species were collected at the same sampling sites except in the southern desert where only HS occurs, thus HB was sampled at 23 different sites and HS was sampled at 30 sites (Fig. 1a,b). At each location, the specific sampling site was selected to maximize the representation of environmental and topographic variation (different slopes, creaks, shade etc.) and minimize anthropogenic disturbance (sampling away from roads, villages, cultivated fields, etc.). Within each sampling site, seeds were collected from 10 different plants with a minimum distance of 10 meters between plants. All HS plants underwent two rounds of single seed descent and selfing, and HB plants underwent only one round of single seed descent due to self-incompatibility. Altogether, 300 HS and 230 HB accessions were collected and analyzed in this study.

Geographic and climatic data for each sampling location were obtained from the Israeli Meteorological Service based on coordinates recorded at each site (Table S1). Soil data was obtained from International Soil Reference and Information Centre (ISRIC) with 1km resolution (Hengl et al., 2014) and internal lab measurements for a soil-sample taken at each site of electric conductivity (EC), field capacity, organic material, inorganic carbon, total carbon, silt and clay content (Table S1).

### Phenotyping under common garden conditions

All seeds from both species were stored in paper bags and oven dried (30°C) for 14 days prior to sowing. After drying, seeds were dipped in 3% chlorine solution for fungal prevention and sown in mid-October in sowing trays, watered, and placed in dark and cold (4°C) conditions for 10 days for vernalization and to break dormancy. After the vernalization period, the trays were transferred to a growing chamber under short day (8h light) conditions at 20°C for germination. After additional 14 days, all accessions were transplanted to 5-liter pots with commercial soil mix (“Ram 158”, Tuff Merom Golan Ltd, Merom Golan, Israel) in a net-house under natural conditions (Hula valley, 33°09’08.4”N 35°37’15.8”E) for the rest of the growing season (Fig. S11).

The phenotyping experiment was conducted in a randomized block design with 4 replicates for each genotype. In HS, the two rounds of selfing and single seed decent enabled to reach a homogenic accessions for replication in a phenotyping experiment. However, the self-incompatibility in HB prevents the fixation of genotypes. To overcome this limitation, one seed was grown until tillering stage which were then cut and split into four pots, thus establishing identical replicates of each genotype.

Phenotypic data was recorded for each plant throughout the growing season for both species and included the following traits: ’days to flowering’ measured from day of transplanting to the emergence of awns with Zadoks scale 49 (Zadoks et al., 1974); ’plant biomass’ measured by weighing all the dry aboveground parts of the plant; ’flag leaf height’ measured from ground level to the base of the highest flag leaf; ’flag leaf length’ measured from the base of the highest flag leaf to its tip; and ’tiller number’ measured at the end of the growing season by counting the number of spike-bearing tillers.

### Measuring genome size

For each accession, ploidy level and genome size were measured using an established flow-cytometry protocol (Doležel et al., 2007; Loureiro et al., 2007). Young leaf tissue was sampled from each genotype in the net-house into liquid nitrogen and taken to the lab for genome size analysis. Each sample was placed in a plastic petri dish along with leaf tissue from maize variety B73, which served as a reference standard with known genome size (Schnable et al., 2009).

Nuclei extraction buffer was prepared by mixing MgCl_2_ (9.15 g L⁻¹), 4-morpholinepropane sulfonate (4.19 g L⁻¹), Sodium Citrate (8.8 g L⁻¹), Sorbitol (21.8 g L⁻¹), Triton X-100, 0.1% w/v (1 mL L⁻¹), L-Dithiothreitol (DTT), and Polyvinylpyrrolidone (1% w/v). The sampled tissues from each genotype together with the maize reference were finely chopped using a disposable razor blade after adding 600 μL of nuclei extraction buffer cooled to 4°C to each petri dish. Once the tissues were homogeneously chopped, they were filtered through 42 μm nylon mesh and transferred into 5 mL tubes. To each tube, 50 μg mL⁻¹ DNA fluorochrome (Propidium Iodide - PI) and 50 μg mL⁻¹ RNase were added and kept on ice with occasional stirring until screening with a flow-cytometer (FACSCalibur™, BD Biosciences Ltd.). In the flow-cytometer, the nuclei sampling rate was adjusted to 20-50 nuclei per second, and a total of 20,000 particles were measured for each sample. The obtained peaks were adjusted in the abscissa using photomultiplier voltage and amplifier gain, while low-channel signals from cell debris and auto-fluorescence were excluded.

The quality of the analysis was evaluated with Flowing Software v.2.5.1 (Turku Centre for Biotechnology, University of Turku, Finland) using the proportion of debris background indicated in the peak distribution symmetry and the coefficient of variation, calculated as the standard deviation of the peak divided by the mean channel position of the peak. This procedure was conducted for two independent tissues from each genotype, and the genome size (2C) was determined in picograms by dividing the product of the reference 2C value (maize) and the sample 2C average by the reference 2C average (Doležel & Bartoš, 2005). Only measurements with peak distribution CV lower than 3% and CV between replicates lower than 2% were considered robust indicators of genome size (Doležel et al., 2007).

All measurements in picograms were converted to base pairs (bp) using the formula: Genome Size (bp) = DNA Content (pg) × 0.978 × 10⁹ (Doležel, J. Barto s. J, Voglmayr H, 2003). As flow-cytometry measures the diploid DNA (2C) genome size, all measurements were divided by ploidy level to obtain values for the haploid level (1C).

### Genotyping the *H. spontaneum* collection

We used whole genome sequencing (WGS) for genotyping the HS collection as described in detail in Potapenko et al. 2025. Briefly, genomic DNA was extracted from a single young seedling using a standard CTAB protocol. Libraries were prepared using the NEBNext® kits (NEB - E7546, NEB - E7595), multiplexed with the Illumina-TruSeq DNA UD Indexes (Illumina - #20022370), and KAPA HiFi HotStart ReadyMix (2X) (KAPABIOSYSTEMS - KK2600). Pooling, titration, and sequencing on the NovaSeq 6000 system.

Raw reads were cleaned trimmed and aligned to the Morex V2 reference genome (Monat et al., 2019) following the GATK4 best practice (Heldenbrand et al., 2019). Variants were filtered with stringent hard filters (Fig. S12) to obtain high-quality SNPs with maximum of 10% missing data across samples and minor allele frequency of 5%. Detailed description of the full procedure and code is provided in Potapenko et al. 2025.

### Genotyping the *H. bulbosum* collection

For each accession, DNA was extracted using the NucleoSpin® Plant II kit (MACHEREY-NAGEL), with cell lysis conducted using the PL2 Buffer. The library preparation and sequencing protocol included DNA digestion of all samples and one negative control with two restriction enzymes, ligation of unique barcoded adapters compatible with the restriction site overhang. Purification and size selection (280-375 bp) were conducted with the Blue Pippin (Sage Science) followed by amplification of libraries via PCR with indexed primers, and finally single-end sequencing on the Illumina NextSeq500.

To identify the best pair of restriction enzymes for *H. bulbosum*, eight double-digested libraries from a pool of 3 samples were tested. The enzyme combinations used were PstI/MspI, PstI/MseI, PstI/NlaIII, PstI/HpyCh4IV, EcoRI/MspI, EcoRI/MseI, EcoRI/NlaIII, and EcoRI/HpyCh4IV. The optimal enzyme combination (EcoRI/NlaIII) was selected to minimize visible repeat regions within the size selection area and maximize amplification. This procedure was conducted by the Australian Genome Research Facility Ltd, Melbourne, Australia.

Raw sequencing reads were demultiplexed, adapter sequences were trimmed, and low-quality reads were removed using the process_radtags module in Stacks v2.0 (Catchen et al., 2013).

Read quality was assessed with FastQC (Andrews, 2017), and high-quality reads were aligned to a haplotype-resolved *Hordeum bulbosum* reference genome (Feng et al., 2025) using BWA-MEM (Li, 2013) with default parameters. Aligned reads were sorted by genomic coordinates with SAMtools v1.9 (Li et al., 2009) and only reads with mapping quality ≥10 were retained for variant calling. Variants were called using the gstacks and populations modules of Stacks, which construct loci from alignments and call genotypes across individuals. Given that the reference genome represents haplotype-resolved sequences the presence of heterozygous genotypes (0/1) is considered an artifact due to sequencing error, misalignment, or unresolved allelic variation. To correct this, we applied a custom Python script that replaced heterozygous genotypes with the homozygous state supported by the higher allele depth or set them to missing if allele depths were equal or ambiguous. Following genotype correction, loci were filtered to retain only high-confidence variants by removing SNPs with >30% missing data and a minor allele frequency (MAF) <5%, using VCFtools (Danecek et al., 2011).

### Statistical analyses of genome size variation

To test for genome size variation between species, the coefficient of variation (Variation = SD/Mean) and quartile coefficient of dispersion (Dispersion = (Q₃-Q₁) / Range) were calculated for each sampling site in each species. These coefficients allow comparison of variation within and between species despite differences in ploidy. In addition, ANOVA was performed to test the differences in genome size variation between sampling sites.

To test for correlations among environment, phenotypic traits and genome size, we conducted a Pearson’s product-moment test between all variables. Correlations between environmental variables and phenotypic traits were assessed using data from sites where both HS and HB co-occur, allowing direct comparison between species. Correlations involving genome size (with either phenotypic traits or environmental variables) were evaluated using the full dataset for each species.

Genome size variation was analyzed using linear mixed-effects models for each species separately, with sampling site included as a random effect to account for non-independence of observations within sites. To test for spatial autocorrelation and dependence, the residuals of non-spatial and spatial (using the sampling locations coordinates) models were tested for autocorrelation using Moran’s I test as implemented in the DHARMa package in R (Hartig, 2024) Prior to model construction, all predictor variables were standardized (mean = 0, SD = 1) to enable direct comparison of effect sizes. Multicollinearity among environmental predictors was assessed using Pearson correlation coefficients, with variables exhibiting correlations >0.8 clustered and reduced to single representatives per cluster. For HS predictor selection retained average annual precipitation (Precipitation), coldest December temperature (December), organic material, total carbon, organic carbon, cation exchange capacity (CECSOL_1km), clay content, sand percentage (SNDPPT_1km), coarse fragments (CRFVOL_1km), electrical conductivity (EC), inorganic content, and inorganic carbon. For HB, 17 predictors were retained, including average annual precipitation (Precipitation), coldest December temperature, soil pH (PHIHOX_1km, PHIKCL_1km), field capacity, organic material, total carbon, organic carbon, cation exchange capacity (CECSOL_1km), clay content, sand content and percentage (SNDPPT_1km), soil pH (PHIHOX_1km, PHIKCL_1km), coarse fragments (CRFVOL_1km), electrical conductivity (EC), inorganic content, and inorganic carbon.

Model selection employed the buildmer R package (Voeten, 2025), which implements forward stepwise selection with cross-validation to determine optimal predictor combinations. The full model scope included all two-way interactions between retained predictors. Final models were fitted using restricted maximum likelihood (REML) estimation. Model assumptions were verified through residual diagnostics (Fig. S4), and variance inflation factors confirmed absence of multicollinearity in final models (VIF < 5). Model performance was evaluated using marginal R² (fixed effects only) and conditional R² (fixed + random effects). Final model for HS included genome size as a function of inorganic carbon, inorganic content, electrical conductivity (EC), soil pH (PHIHOX_1km), precipitation, and the precipitation × coarse fragments interaction, with sampling site as a random intercept to account for non-independence of individuals belonging to the same site. Final model for HB was genome size as a function of cation exchange capacity (CECSOL_1km), coarse fragments (CRFVOL_1km), precipitation × coarse fragments interaction, clay × organic material interaction, and organic material × field capacity interaction, with sampling site information as a random intercept.

### Population genomics analyses

Population stratification and diversity analyses were conducted for each species separately using the variants called in each species (WGS or GBS). Each SNP dataset was LD-pruned with a minimum LD cut-off of r² = 0.2 before conducting principal component analysis (PCA) using the LEA package in R (Frichot & François, 2015). To further investigate population stratification, structure analysis was performed on LD-prunned datasets using the sNMF approach implemented in LEA. This method uses multilocus genotype data to identify the number of populations in a dataset and assign individuals to each inferred population. Population differentiation was assessed using Analysis of Molecular Variance (AMOVA) to test the genetic variation among populations (Fig. S1). Variant data in VCF format were first filtered to a random subset of 10,000 SNPs to reduce computational load. The VCF file was loaded into R using the vcfR package (Knaus & Grünwald, 2017). Pairwise genetic distances among individuals were calculated using bitwise Euclidean distances from poppr v2.9.6 package (Kamvar et al., 2015).

AMOVA was performed with the poppr.amova function using the calculated distance matrix, testing the partitioning of genetic variance between and within populations. The significance of the observed variance components was assessed with a permutation test (randtest) using 999 random permutations.

### Quantifying repetitive elements

Quantifying the abundance of repetitive elements along the genome was conducted only for HS where whole genome sequence data is available. We used the dnaPipeTE software v1.3.1(Goubert et al., 2015) which allows finding, annotating, and quantifying repetitive elements in a representative fraction of whole genome sequencing data. To screen for repetitive element sequences, a reference panel was established using all sequences tagged as “Hordeum” in the Triticeae Repetitive Elements Platform (TREP) non redundant database v19 (Wicker et al., 2002). To avoid sampling bias in repetitive elements frequencies, an equal number of random subsets of reads was sampled for each individual and used in the analysis.

To identify the minimum number of reads required to properly represent the genome composition, the analysis was performed with varying subsets of reads from 100,000 to 10,000,000, obtained from one individual, and the proportion of repetitive elements identified was evaluated at each round. Once the proportion of different repetitive element types has remained constant along exceeding number of reads, sampling was saturated. The minimum number of reads to reach saturation was 1,000,000, which was fixed as the number of reads sampled across all individuals. The actual number of repetitive elements in base-pairs was then calculated for the entire genome based on the subset of 1,000,000 reads of 150 bp. To achieve this, the fraction of repetitive elements was multiplied by a factor (Genome Size (bp) / (No. of Reads * Read Length)).

### Calculating divergence among transposable elements

To investigate the temporal dynamics of transposable element activity, we first annotated repeats using RepeatMasker v4.1.5 (Smit et al., 2013) and the non-redundant TREP database (Wicker et al., 2002) after extracting sequences specific to *Hordeum* species. As a reference genome for HS, we used a high-quality genome assembly B1K-04-12 V1 (Jayakodi et al., 2020b), a wild barley accession that was sampled in Israel (Hübner et al., 2009).

Following TE annotation, we estimated the age of repeat insertions using the RepeatMasker utility script calcDivergenceFromAlign.pl, which computes sequence divergence between TE copies and their corresponding consensus sequences in the database. This script applies the Kimura 2-parameter (K2P) model to account for multiple substitutions at the same site and transitions versus transversions. The resulting divergence values reflect the accumulated mutations since the insertion of each TE. To convert divergence values to approximate insertion time, we applied a standard substitution rate widely used in plant repeat studies: *T = K / (2 * r)* where *T* is the estimated time since insertion (in million years), *K* is the Kimura-corrected divergence, *r* is the neutral substitution rate, set to 1.3x10^-8^ substitutions per site per year (Bowen & McDonald, 2001; Ma & Bennetzen, 2004). The resulting age distribution of TE insertions was then used to infer historical bursts of transposition and to compare repeat dynamics between genomes.

### Genome-wide association studies

Genome-wide association studies (GWAS) were performed for the full SNP datasets using the EMMAX software (Kang et al., 2010). For correction of population structure, the first three principal components obtained from the PCA of the LD-pruned SNP dataset (described above) were used, and genetic relatedness was corrected based on a Balding-Nichols kinship matrix (Balding & Nichols, 1995). Phenotype data was converted to BLUP scores using a mixed linear model implemented in the lme4 package (Bates et al., 2015). In these models, genotypes, season, and block number were included as random effects to account for experimental design and environmental variation. The maximum number of iterations was increased to 10,000 to ensure proper model convergence. Due to the stringent Bonferroni correction, a high false negative rate was observed, genomic regions were considered significantly associated when at least one SNP exceeded the P < 1 × 10⁻⁶ threshold for HS and P < 1 x 10^-4^ threshold for HB. The boundaries of this region were defined by the extent of contiguous association signals above P < 1 × 10⁻⁵. Candidate genes located within the defined region were identified based on the reference genome annotation. For genes with uncertain or putative annotations, we conducted protein sequence similarity searches using the NCBI BLASTP web service with default parameters (Johnson et al., 2008). The visualization of GWAS results were obtained using the CMplot package (Yin et al., 2021) and the topr R package (Juliusdottir, 2023).

Following GWAS, SNPs within the genomic region on chromosome 3H associated with genome size in HS were extracted and thinned using VCFtools at 250 kb intervals, randomly selecting one SNP per interval to reduce marker density for clustering. Local haplotype structure was inferred with the R package CrossHap (Marsh et al., 2023), an LD-based haplotyping tool for visualizing patterns of variation across loci and individuals. Each haplotype was then compared to the adaptive haplotype using Wilcoxon rank-sum tests for genome size.

### Identifying selection sweeps

We applied the RAiSD algorithm (Alachiotis & Pavlidis, 2018) to detect signatures of selective sweeps among individuals carrying the different haplotypes around the genomic region associated with genome size variation. RAiSD calculates a composite μ statistic from the site frequency spectrum, linkage disequilibrium, and changes in diversity across empirically defined windows of 200 SNPs. To further support the selective sweep detection, we calculated independently the nucleotide diversity (π) and Tajima’s D in sliding windows of 3Mbp along the genome using VCFtools (Danecek et al., 2011).

## References

Alachiotis, N., & Pavlidis, P. (2018). RAiSD detects positive selection based on multiple signatures of a selective sweep and SNP vectors. Communications Biology 2018 1:1, 1(1), 1–11. 10.1038/s42003-018-0085-8

Ammari, M., Maseh, K., & Zander, M. (2024). PIF transcription factors-versatile plant epigenome landscapers. Frontiers in Epigenetics and Epigenomics, 2, 1404958. 10.3389/FREAE.2024.1404958

Ammiraju, J. S. S., Zuccolo, A., Yu, Y., Song, X., Piegu, B., Chevalier, F., Walling, J. G., Ma, J., Talag, J., Brar, D. S., Sanmiguel, P. J., Jiang, N., Jackson, S. A., Panaud, O., & Wing, R. A. (2007). Evolutionary dynamics of an ancient retrotransposon family provides insights into evolution of genome size in the genus Oryza. Wiley Online Library, 52(2), 342–351. 10.1111/j.1365-313X.2007.03242.x

Andrews, S. (2017). FastQC: a quality control tool for high throughput sequence data.2010.

Arumuganathan, K., & Earle, E. D. (1991). Nuclear DNA content of some important plant species. Plant Molecular Biology Reporter, 9(3), 208–218. 10.1007/BF02672069

Baduel, P., Quadrana, L., Hunter, B., Bomblies, K., & Colot, V. (2019). Relaxed purifying selection in autopolyploids drives transposable element over-accumulation which provides variants for local adaptation. Nature Communications 2019 10:1, 10(1), 1–10. 10.1038/s41467-019-13730-0

Balding, D. J., & Nichols, R. A. (1995). A method for quantifying differentiation between populations at multi-allelic loci and its implications for investigating identity and paternity. Genetica, 96(1–2), 3–12. 10.1007/BF01441146/METRICS

Basak, S., Sun, X., Wang, G., & Yang, Y. (2019). Genome Size Unaffected by Variation in Morphological Traits, Temperature, and Precipitation in Turnip. Applied Sciences 2019, Vol. 9, Page *253*, 9(2), 253. 10.3390/APP9020253

Bates, D., Mächler, M., Bolker, B. M., & Walker, S. C. (2015). Fitting Linear Mixed-Effects Models Using lme4. Journal of Statistical Software, 67(1), 1–48. 10.18637/JSS.V067.I01

Beaulieu, J. M., Leitch, I. J., Patel, S., Pendharkar, A., & Knight, C. A. (2008). Genome size is a strong predictor of cell size and stomatal density in angiosperms. Wiley Online Library, 179(4), 975–986. 10.1111/j.1469-8137.2008.02528.x

Beaulieu, J. M., Moles, A. T., Leitch, I. J., Bennett, M. D., Dickie, J. B., & Knight, C. A. (2007). Correlated evolution of genome size and seed mass. New Phytologist, 173(2), 422. 10.1111/j.1469-8137.2006.01919.x

Bennett, M. D., & Leitch, I. J. (2005). Genome Size Evolution in Plants. The Evolution of the Genome, 89–162. 10.1016/B978-012301463-4/50004-8

Benor, S., Fuchs, J., Blattner, F. R., & Fuchs, J. (2011). Genome size variation in Corchorus olitorius (Malvaceae sl) and its correlation with elevation and phenotypic traits. Cdnsciencepub.Com, 54(7), 575–585. 10.1139/g11-021

Bilinski, P., Albert, P. S., Berg, J. J., Birchler, J. A., Grote, M. N., Lorant, A., Quezada, J., Swarts, K., Yang, J., & Ross-Ibarra, J. (2018). Parallel altitudinal clines reveal trends in adaptive evolution of genome size in Zea mays. PLOS Genetics, 14(5), e1007162. 10.1371/JOURNAL.PGEN.1007162

Blattner, F. R. (2006). Multiple intercontinental dispersals shaped the distribution area of Hordeum (Poaceae). New Phytologist, 169(3), 603–614. 10.1111/J.1469-8137.2005.01610.X

Blattner, F. R. (2018). Taxonomy of the Genus Hordeum and Barley (Hordeum vulgare). 11–23. 10.1007/978-3-319-92528-8_2

Bothmer, R., Jacobsen, N., Baden, C., & Jorgensen, R. (1995). An ecogeographical study of the genus Hordeum. https://cgspace.cgiar.org/handle/10568/104311

Bourque, G., Burns, K. H., Gehring, M., Gorbunova, V., Seluanov, A., Hammell, M., Imbeault, M., Izsvák, Z., Levin, H. L., Macfarlan, T. S., Mager, D. L., & Feschotte, C. (2018). Ten things you should know about transposable elements 06 Biological Sciences 0604 Genetics. Genome Biology, 19(1). 10.1186/S13059-018-1577-Z

Bowen, N. J., & McDonald, J. F. (2001). Drosophila Euchromatic LTR Retrotransposons are Much Younger Than the Host Species in Which They Reside. Genome Research, 11(9), 1527–1540. 10.1101/GR.164201

Catchen, J., Hohenlohe, P. A., Bassham, S., Amores, A., & Cresko, W. A. (2013). Stacks: An analysis tool set for population genomics. Molecular Ecology, 22(11), 3124–3140. 10.1111/MEC.12354

Cavalier-Smith, T. (1985). Eukaryote Gene Numbers, Non Coding DNA and Genome Size. The Evolution of Genome Size, 69–103. https://cir.nii.ac.jp/crid/1571980074173088384

Danecek, P., Auton, A., Abecasis, G., Albers, C. A., Banks, E., DePristo, M. A., Handsaker, R. E., Lunter, G., Marth, G. T., Sherry, S. T., McVean, G., & Durbin, R. (2011). The variant call format and VCFtools. Bioinformatics, 27(15), 2156–2158. 10.1093/BIOINFORMATICS/BTR330

Doležel, J. Barto s. J, Voglmayr H, G. J. (2003). Nuclear DNA content and genome size of trout and human. *Cytometry*, A, 51(2), 127–128.

Doležel, J., & Bartoš, J. (2005). Plant DNA Flow Cytometry and Estimation of Nuclear Genome Size. Annals of Botany, 95(1), 99–110. 10.1093/AOB/MCI005

Doležel, J., Greilhuber, J., protocols, J. S.-N., & 2007, undefined. (2007). Estimation of nuclear DNA content in plants using flow cytometry. Nature.Com, 2(9), 2233–2244. 10.1038/nprot.2007.310

Du, Y. P., Bi, Y., Zhang, M. F., Yang, F. P., Jia, G. X., & Zhang, X. H. (2017). Genome size diversity in Lilium (Liliaceae) is correlated with karyotype and environmental traits. Frontiers in Plant Science, 8. 10.3389/FPLS.2017.01303/FULL

Durán-Figueroa, N., & Vielle-Calzada, J. P. (2010). ARGONAUTE9-dependent silencing of transposable elements in pericentromeric regions of Arabidopsis. Plant Signaling and Behavior, 5(11). 10.4161/PSB.5.11.13548/ASSET/D7D0E576-EB4D-4766-B484-2D314399AD40/ASSETS/IMAGES/LARGE/KPSB_A_10913548_F0002.JPG

Esnault, C., Lee, M., Ham, C., & Levin, H. L. (2019). Transposable element insertions in fission yeast drive adaptation to environmental stress. Genome Research, 29(1), 85–95. 10.1101/GR.239699.118/-/DC1

Feng, J. W., Pidon, H., Cuacos, M., Lux, T., Himmelbach, A., Haghi, R., Fuchs, J., Haberer, G., Kuo, Y. T., Guo, Y., Jayakodi, M., Toegelová, H., Harpke, D., Knauft, M., Fiebig, A., Maruschewski, M., Ronen, M., Sharon, A., Šimková, H., … Mascher, M. (2025). A haplotype-resolved pangenome of the barley wild relative Hordeum bulbosum. Nature 2025, 1–10. 10.1038/s41586-025-09270-x

Feschotte, C., & Pritham, E. J. (2007). DNA transposons and the evolution of eukaryotic genomes. Annual Review of Genetics, 41, 331–368. 10.1146/ANNUREV.GENET.40.110405.090448

Frichot, E., & François, O. (2015). LEA: An R package for landscape and ecological association studies. Methods in Ecology and Evolution, 6(8), 925–929. 10.1111/2041-210X.12382

Fuerst, D., Shermeister, B., Mandel, T., & Hübner, S. (2023). Evolutionary Conservation and Transcriptome Analyses Attribute Perenniality and Flowering to Day-Length Responsive Genes in Bulbous Barley (Hordeum bulbosum). Genome Biology and Evolution, 15(1). 10.1093/GBE/EVAC168

Gallagher, R. V, Leishman, M. R., Miller, J. T., Hui, C., Richardson, D. M., Suda, J., & Trávníček, P. (2011). Invasiveness in introduced Australian acacias: the role of species traits and genome size. Wiley Online Library, 17(5), 884–897. 10.1111/j.1472-4642.2011.00805.x

Goubert, C., Modolo, L., Vieira, C., Moro, C. V., Mavingui, P., & Boulesteix, M. (2015). De novo assembly and annotation of the Asian tiger mosquito (Aedesalbopictus) repeatome with dnaPipeTE from raw genomic reads and comparative analysis with the yellow fever mosquito (Aedes aegypti). Genome Biology and Evolution, 7(4), 1192– 1205. 10.1093/GBE/EVV050

Graham, M. J., Nickell, C. D., & Rayburn, A. L. (1994). Relationship between genome size and maturity group in soybean. Theoretical and Applied Genetics, 88(3–4), 429–432. 10.1007/BF00223656

Gregory, T. R. (2001). Coincidence, coevolution, or causation? DNA content, cell size, and the C-value enigma. Biological Reviews of the Cambridge Philosophical Society, 76(1), 65–101. 10.1017/S1464793100005595

Hartig, F. (2024). DHARMa: Residual Diagnostics for Hierarchical (Multi-Level / Mixed) Regression Models. https://github.com/florianhartig/dharma

Heldenbrand, J. R., Baheti, S., Bockol, M. A., Drucker, T. M., Hart, S. N., Hudson, M. E., Iyer, R. K., Kalmbach, M. T., Kendig, K. I., Klee, E. W., Mattson, N. R., Wieben, E. D., Wiepert, M., Wildman, D. E., & Mainzer, L. S. (2019). Recommendations for performance optimizations when using GATK3.8 and GATK4. BMC Bioinformatics, 20(1), 1–9. 10.1186/S12859-019-3169-7/TABLES/4

Hengl, T., De Jesus, J. M., MacMillan, R. A., Batjes, N. H., Heuvelink, G. B. M., Ribeiro, E., Samuel-Rosa, A., Kempen, B., Leenaars, J. G. B., Walsh, M. G., & Gonzalez, M. R. (2014). SoilGrids1km — Global Soil Information Based on Automated Mapping. PLOS ONE, 9(8), e105992. 10.1371/JOURNAL.PONE.0105992

Herben, T., Suda, J., Klimešová, J., Mihulka, S., Íha, P., & Šímová, I. (2012). Ecological effects of cell-level processes: genome size, functional traits and regional abundance of herbaceous plant species. Annals of Botany, 110(7), 1357–1367. 10.1093/AOB/MCS099

Hübner, S., Bdolach, E., … S. E.-J. of, & 2013, undefined. (2013). Phenotypic landscapes: phenological patterns in wild and cultivated barley. Wiley Online Library, 26(1), 163–174. 10.1111/jeb.12043

Hübner, S., Günther, T., Flavell, A., Fridman, E., Graner, A., Korol, A., & Schmid, K. J. (2012). Islands and streams: Clusters and gene flow in wild barley populations from the Levant. Molecular Ecology, 21(5), 1115–1129. 10.1111/J.1365-294X.2011.05434.X

Hübner, S., HÖffken, M., Oren, E., Haseneyer, G., Stein, N., Graner, A., Schmid, K., & Fridman, E. (2009). Strong correlation of wild barley (Hordeum spontaneum) population structure with temperature and precipitation variation. Molecular Ecology, 18(7), 1523–1536. 10.1111/J.1365-294X.2009.04106.X

Hübner, S., & Kantar, M. B. (2021). Tapping Diversity From the Wild: From Sampling to Implementation. Frontiers in Plant Science, 12. 10.3389/FPLS.2021.626565/FULL

Jakob, S. S., Meister, A., & Blattner, F. R. (2004). The Considerable Genome Size Variation of Hordeum Species (Poaceae) Is Linked to Phylogeny, Life Form, Ecology, and Speciation Rates. Molecular Biology and Evolution, 21(5), 860–869. 10.1093/MOLBEV/MSH092

Jayakodi, M., Padmarasu, S., Haberer, G., Bonthala, V. S., Gundlach, H., Monat, C., Lux, T., Kamal, N., Lang, D., Himmelbach, A., Ens, J., Zhang, X. Q., Angessa, T. T., Zhou, G., Tan, C., Hill, C., Wang, P., Schreiber, M., Boston, L. B., … Stein, N. (2020a). The barley pan-genome reveals the hidden legacy of mutation breeding. Nature 2020 588:7837, 588(7837), 284–289. 10.1038/s41586-020-2947-8

Jayakodi, M., Padmarasu, S., Haberer, G., Bonthala, V. S., Gundlach, H., Monat, C., Lux, T., Kamal, N., Lang, D., Himmelbach, A., Ens, J., Zhang, X. Q., Angessa, T. T., Zhou, G., Tan, C., Hill, C., Wang, P., Schreiber, M., Boston, L. B., … Stein, N. (2020b). The barley pan-genome reveals the hidden legacy of mutation breeding. Nature 2020 588:7837, 588(7837), 284–289. 10.1038/s41586-020-2947-8

Johnson, M., Zaretskaya, I., Raytselis, Y., Merezhuk, Y., McGinnis, S., & Madden, T. L. (2008). NCBI BLAST: a better web interface. Nucleic Acids Research, 36(suppl_2), W5– W9. 10.1093/NAR/GKN201

Jordan, G. J., Carpenter, R. J., Koutoulis, A., Price, A., & Brodribb, T. J. (2015). Environmental adaptation in stomatal size independent of the effects of genome size. New Phytologist, 205(2), 608–617. 10.1111/NPH.13076

Jörgensen, R. B. (1982). Biosystematics of Hordeum bulbosum L. Nordic Journal of Botany, 2(5), 421–434. 10.1111/J.1756-1051.1982.TB01205.X

Juliusdottir, T. (2023). topr: an R package for viewing and annotating genetic association results. BMC Bioinformatics, 24(1), 1–10. 10.1186/S12859-023-05301-4/FIGURES/3

Kamvar, Z. N., Brooks, J. C., & Grünwald, N. J. (2015). Novel R tools for analysis of genome-wide population genetic data with emphasis on clonality. Frontiers in Genetics, 6(JUN), 151034. 10.3389/FGENE.2015.00208/BIBTEX

Kang, H. M., Sul, J. H., Service, S. K., Zaitlen, N. A., Kong, S. Y., Freimer, N. B., Sabatti, C., & Eskin, E. (2010). Variance component model to account for sample structure in genome-wide association studies. Nature Genetics 2010 42:4, 42(4), 348–354. 10.1038/ng.548

Kankanpaa, J., Mannonen, L., & Schulman, A. H. (1996). The genome sizes of Hordeum species show considerable variation. Genome, 39(4), 730–735. 10.1139/G96-092

Khorami, S. S., Arzani, K., Karimzadeh, G., Shojaeiyan, A., & Ligterink, W. (2018). Genome Size: A Novel Predictor of Nut Weight and Nut Size of Walnut Trees. HortScience, 53(3), 275–282. 10.21273/HORTSCI12725-17

Knaus, B. J., & Grünwald, N. J. (2017). vcfr: a package to manipulate and visualize variant call format data in R. Molecular Ecology Resources, 17(1), 44–53. 10.1111/1755-0998.12549

Knight, C. A., Molinari, N. A., & Petrov, D. A. (2005). The Large Genome Constraint Hypothesis: Evolution, Ecology and Phenotype. Annals of Botany, 95(1), 177–190. 10.1093/AOB/MCI011

Koonin, E. V. (2016). Splendor and misery of adaptation, or the importance of neutral null for understanding evolution. BMC Biology, 14(1), 1–8. 10.1186/S12915-016-0338-2/FIGURES/4

Krahulcová, A., Trávníček, P., Krahulec, F., & Rejmánek, M. (2017). Small genomes and large seeds: chromosome numbers, genome size and seed mass in diploid Aesculus species (Sapindaceae). Annals of Botany, 119(6), 957–964. 10.1093/AOB/MCW261

LaBar, T., & Adami, C. (2020). Genome Size and the Extinction of Small Populations. 167–183. 10.1007/978-3-030-39831-6_14

Lee, J., Waminal, N. E., Choi, H. Il, Perumal, S., Lee, S. C., Nguyen, V. B., Jang, W., Kim, N. H., Gao, L. Z., & Yang, T. J. (2017). Rapid amplification of four retrotransposon families promoted speciation and genome size expansion in the genus Panax. Scientific Reports 2017 7:1, 7(1), 1–9. 10.1038/s41598-017-08194-5

Li, H. (2013). Aligning sequence reads, clone sequences and assembly contigs with BWA- MEM. http://arxiv.org/abs/1303.3997

Li, H., Handsaker, B., Wysoker, A., Fennell, T., Ruan, J., Homer, N., Marth, G., Abecasis, G., & Durbin, R. (2009). The Sequence Alignment/Map format and SAMtools. Bioinformatics, 25(16), 2078–2079. 10.1093/BIOINFORMATICS/BTP352

Liang, S. C., Hartwig, B., Perera, P., Mora-García, S., de Leau, E., Thornton, H., de Alves, F. L., Rapsilber, J., Yang, S., James, G. V., Schneeberger, K., Finnegan, E. J., Turck, F., & Goodrich, J. (2015). Kicking against the PRCs – A Domesticated Transposase Antagonises Silencing Mediated by Polycomb Group Proteins and Is an Accessory Component of Polycomb Repressive Complex 2. PLOS Genetics, 11(12), e1005660. 10.1371/JOURNAL.PGEN.1005660

Loureiro, J., Rodriguez, E., Doležel, J., & Santos, C. (2007). Two New Nuclear Isolation Buffers for Plant DNA Flow Cytometry: A Test with 37 Species. Annals of Botany, 100(4), 875–888. 10.1093/AOB/MCM152

Lynch, M., & Conery, J. S. (2003). The Origins of Genome Complexity. Science, 302(5649), 1401–1404. 10.1126/science.1089370

Ma, J., & Bennetzen, J. L. (2004). Rapid recent growth and divergence of rice nuclear genomes. Proceedings of the National Academy of Sciences of the United States of America, 101(34), 12404–12410. 10.1073/PNAS.0403715101/SUPPL_FILE/03715TABLE6.HTML

Marsh, J. I., Petereit, J., Johnston, B. A., Bayer, P. E., Tay Fernandez, C. G., Al-Mamun, H. A., Batley, J., & Edwards, D. (2023). crosshap: R package for local haplotype visualization for trait association analysis. Bioinformatics, 39(8). 10.1093/BIOINFORMATICS/BTAD518

Mei, W., Stetter, M. G., Gates, D. J., Stitzer, M. C., & Ross-Ibarra, J. (2018). Adaptation in plant genomes: Bigger is different. American Journal of Botany, 105(1), 16–19. 10.1002/ajb2.1002

Michaelson, M. J., Price, H. J., Johnston, J. S., & Ellison, J. R. (1991). VARIATION OF NUCLEAR DNA CONTENT IN HELIANTHUS ANNUUS (ASTERACEAE). American Journal of Botany, 78(9), 1238–1243. 10.1002/J.1537-2197.1991.TB11416.X

Mirsky, A. E., & Ris, H. (1951). The desoxyribonucleic acid content of animal cells and its evolutionary significance. The Journal of General Physiology, 34(4), 451. 10.1085/JGP.34.4.451

Mitsuda, N., Seki, M., Shinozaki, K., & Ohme-Takagi, M. (2005). The NAC Transcription Factors NST1 and NST2 of Arabidopsis Regulate Secondary Wall Thickenings and Are Required for Anther Dehiscence. The Plant Cell, 17(11), 2993. 10.1105/TPC.105.036004

Monat, C., Padmarasu, S., Lux, T., Wicker, T., Gundlach, H., Himmelbach, A., Ens, J., Li, C., Muehlbauer, G. J., Schulman, A. H., Waugh, R., Braumann, I., Pozniak, C., Scholz, U., Mayer, K. F. X., Spannagl, M., Stein, N., & Mascher, M. (2019). TRITEX: Chromosome-scale sequence assembly of Triticeae genomes with open-source tools. Genome Biology, 20(1), 1–18. 10.1186/S13059-019-1899-5/FIGURES/6

Morgan, H. D., & Westoby, M. (2005). The Relationship Between Nuclear DNA Content and Leaf Strategy in Seed Plants. Annals of Botany, 96(7), 1321–1330. 10.1093/AOB/MCI284

Petrov, D. A. (2001). Evolution of genome size: New approaches to an old problem. Trends in Genetics, 17(1), 23–28. 10.1016/S0168-9525(00)02157-0

Piegu, B., Guyot, R., Picault, N., Roulin, A., Saniyal, A., Kim, H., Collura, K., Brar, D. S., Jackson, S., Wing, R. A., & Panaud, O. (2006). Doubling genome size without polyploidization: Dynamics of retrotransposition-driven genomic expansions in Oryza australiensis, a wild relative of rice. Genome Research, 16(10), 1262–1269. 10.1101/GR.5290206

Potapenko, E. V., Schwarts, D., Mandel, T., Ashkenazy, N., Fuerst, D., Atsmon, G., Korol, A. B., Kantar, M. B., Bar-Massada, A., & Hübner, S. (2025). Not so selfish: transposable elements fuel adaptation to climate change. BioRxiv, 2024.09.02.610836. 10.1101/2024.09.02.610836

Rayburn, A. L., & Auger, J. A. (1990). Genome size variation in Zea mays ssp. mays adapted to different altitudes. Theoretical and Applied Genetics, 79(4), 470–474. 10.1007/BF00226155

Rayburn, A. L., Birdar, D. P., Bullock, D. G., Nelson, R. L., Gourmet, C., & Wetzel, J. B. (1997). Nuclear DNA content diversity in Chinese soybean introductions. Elsevier, 80, 321–325. https://www.sciencedirect.com/science/article/pii/S0305736497904455

Realini, M. F., Poggio, L., Cámara-Hernández, J., & González, G. E. (2016). Intra-specific variation in genome size in maize: cytological and phenotypic correlates. AoB PLANTS, 8. 10.1093/AOBPLA/PLV138

Reeves, G., Francist, D., Davies, M. S., Rogers, H. J., & Hodkinsont, T. R. (1998). Genome Size is Negatively Correlated with Altitude in Natural Populations of Dactylis glomerata. Academic.Oup.Com, 82, 99–105. https://academic.oup.com/aob/article-abstract/82/suppl_1/99/211227

Roddy, A. B., Théroux-Rancourt, G., Abbo, T., Benedetti, J. W., Brodersen, C. R., Castro, M., Castro, S., Gilbride, A. B., Jensen, B., Jiang, G. F., Perkins, J. A., Perkins, S. D., Loureiro, J., Syed, Z., Alexander Thompson, R., Kuebbing, S. E., & Simonin, K. A. (2020). The Scaling of Genome Size and Cell Size Limits Maximum Rates of Photosynthesis with Implications for Ecological Strategies. 10.1086/706186, 181(1), 75–87. https://doi.org/10.1086/706186

Schnable, P. S., Ware, D., Fulton, R. S., Stein, J. C., Wei, F., Pasternak, S., Liang, C., Zhang, J., Fulton, L., Graves, T. A., Minx, P., Reily, A. D., Courtney, L., Kruchowski, S. S., Tomlinson, C., Strong, C., Delehaunty, K., Fronick, C., Courtney, B., … Wilson, R. K. (2009). The B73 maize genome: Complexity, diversity, and dynamics. Science, 326(5956), 1112–1115. 10.1126/SCIENCE.1178534

Schubert, I., science, G. V.-T. in plant, & 2016, undefined. (2016). Genome stability and evolution: attempting a holistic view. Elsevier. https://www.sciencedirect.com/science/article/pii/S1360138516300681?casa_token=BlhVTkuZNjIAAAAA:G_gdMj1gFjRHJZJ47Rjx-BxmV4BuXmG37N5desieIUgjCOeSj_45rKZXGDYyjNbn4InD_A

Smit, A. F. A., Hubley, R., & Green, P. (2013). RepeatMasker Open-4.0.

Stapley, J., Santure, A. W., & Dennis, S. R. (2015). Transposable elements as agents of rapid adaptation may explain the genetic paradox of invasive species. Molecular Ecology, 24(9), 2241–2252. 10.1111/MEC.13089

Turpeinen, T., Kulmala, J., & Nevo, E. (1999). Genome size variation in Hordeum spontaneum populations. Genome, 42(6), 1094–1099. 10.1139/G99-066

Underwood, C. J., & Martienssen, R. A. (2015). Argonautes team up to silence transposable elements in Arabidopsis. The EMBO Journal, 34(5), 579. 10.15252/EMBJ.201590971

Voeten, C. C. (2025). buildmer: Stepwise Elimination and Term Reordering for Mixed-Effects Regression.

von Bothmer, R., & Jacobsen, N. (2015). Origin, Taxonomy, and Related Species. 19–56. 10.2134/AGRONMONOGR26.C2

Wang, L., Li, H., Lei, Z., Jeong, D. H., & Cho, J. (2024). The CARBON CATABOLITE REPRESSION 4A-mediated RNA deadenylation pathway acts on the transposon RNAs that are not regulated by small RNAs. New Phytologist, 241(4), 1636–1645. 10.1111/NPH.19435

Whitney, K. D., Baack, E. J., Hamrick, J. L., Godt, M. J. W., Barringer, B. C., Bennett, M. D., Eckert, C. G., Goodwillie, C., Kalisz, S., Leitch, I. J., & Ross-Ibarra, J. (2010). A role for nonadaptive processes in plant genome size evolution? Evolution, 64(7), 2097–2109. 10.1111/j.1558-5646.2010.00967.x

Wicker, T., Matthews, D., & Keller, B. (2002). TREP: a database for Triticeae repetitive elements. http://botserv2.uzh.ch/kelldata/trep-db/pdfs/2002_TIPS.pdf

Wicker, T., Schulman, A. H., Tanskanen, J., Spannagl, M., Twardziok, S., Mascher, M., Springer, N. M., Li, Q., Waugh, R., Li, C., Zhang, G., Stein, N., Mayer, K. F. X., & Gundlach, H. (2017). The repetitive landscape of the 5100 Mbp barley genome. Mobile DNA, 8(1). 10.1186/S13100-017-0102-3

Wu-Scharf, D., Jeong, B. R., Zhang, C., & Cerutti, H. (2000). Transgene and Transposon Silencing in Chlamydomonas reinhardtii by a DEAH-Box RNA Helicase. Science, 290(5494), 1159–1162. 10.1126/SCIENCE.290.5494.1159

Yin, L., Zhang, H., Tang, Z., Xu, J., Yin, D., Zhang, Z., Yuan, X., Zhu, M., Zhao, S., Li, X., & Liu, X. (2021). rMVP: A Memory-Efficient, Visualization-Enhanced, and Parallel-Accelerated Tool for Genome-Wide Association Study. *Genomics*, Proteomics & Bioinformatics, 19(4), 619–628. 10.1016/J.GPB.2020.10.007

Zadoks, J. C., Chang, T. T., & Konzak, C. F. (1974). A decimal code for the growth stages of cereals. 14, 415–421.

Zhang, Q., Wang, Z., Lu, X., Yan, H., Zhang, H., He, H., Bischof, S., Jake Harris, C., & Liu, Q. (2023). DDT-RELATED PROTEIN4–IMITATION SWITCH alters nucleosome distribution to relieve transcriptional silencing in Arabidopsis. The Plant Cell, 35(8), 3109–3126. 10.1093/PLCELL/KOAD143

Zhong, J., He, W., Peng, Z., Zhang, H., Li, F., & Yao, J. (2019). A putative AGO protein, OsAGO17, positively regulates grain size and grain weight through OsmiR397b in rice. Plant Biotechnology Journal, 18(4), 916. 10.1111/PBI.13256

